# The changing morphology of the ventricular walls of mouse and human with increasing gestation

**DOI:** 10.1101/2023.11.05.565685

**Authors:** Bjarke Jensen, Yun Hee Chang, Simon D. Bamforth, Timothy Mohun, David Sedmera, Martin Bartos, Robert H. Anderson

## Abstract

That the highly trabeculated ventricular walls of the developing embryos transforms to the arrangement during the fetal stages, when the mural architecture is dominated by the thickness of the compact myocardium, has been explained by coalescence of trabeculations, often erroneously described as “compaction”. Recent data, however, supports differential rates of growth of the trabecular and compact layers as the major driver of change. Here, these processes were assessed quantitatively and morphologically using a larger dataset than has previously been available of mouse hearts from embryonic day 10.5 to postnatal day 3, supported by images from human hearts. The volume of the trabecular layer increased throughout development, in contrast to what would be expected had there been “compaction”. During the embryonic-fetal transition, fast growth of the compact layer diminished the proportion of trabeculations. Similarly, great expansion of the central cavity reduced the proportion that intertrabecular recesses make of the total cavity. Using the median value of left ventricular trabeculation, we provided illustrations for each gestational day so as to provide pictorial evidence of the changes. The illustrations confirmed a pronounced growth of the compact wall, and prominence of the central cavity. This corresponds, in morphological terms, to a reduction in the extent of the trabecular layer. Similar observations were made in the human hearts. We conclude that it is a period of comparatively slow growth of the trabecular layer, rather than so-called compaction, that is the major determinant of the changing morphology of the ventricular walls of both mouse and human.

## Introduction

Differential growth rates have long been viewed as a major transformative factor of morphological change, both in ontogeny and evolution (Gould, 1966). In humans, the tremendous growth involved in transforming the fertilized egg into a neonate weighing approximately 3.5 kilograms allows for a great scope for such differential rates to impact on morphology, including the proportions of the organs and their size (de Bakker et al., 2016). For the heart, much of the transformation from the stage of a tube with a solitary lumen to the 4-chambered organ can be attributed to the rapid growth of the chamber walls, which ‘balloon’ out from the slow-growing primary myocardium that constitutes the walls of the heart tube (Moorman and Christoffels, 2003). This growth is driven by cardiomyocytic proliferation, which itself is governed by evolutionarily-conserved transcriptional networks (Jensen et al., 2013, Sedmera et al., 2003, de Boer et al., 2012, Tian et al., 2017, Wang et al., 2018). Recently, using developmental series of human, mouse, shrew, and chicken hearts, differential growth rates were shown to explain much of the gestational change found in the ratio of the width of the trabecular and compact components of the ventricular walls (Faber et al., 2022b, ChangSheftel and Jensen, 2022). The ratio of trabecular-to-compact myocardium is important in so far that it is a widely used metric for diagnosing so-called left ventricular non-compaction cardiomyopathy, notwithstanding that the diagnosis of this alleged entity has become controversial because of a very substantial rate of overdiagnosis (Ross et al., 2020, Anderson et al., 2017). In fact, the mere presence of excessive trabeculation can be considered to be without prognostic value (Aung et al., 2020, Petersen et al., 2023).

Non-compaction should be understood as the absence of compaction (Chin et al., 1990). Compaction itself is a process to which much importance has been assigned in transforming the highly trabeculated embryonic cardiac ventricular walls to the walls as seen in fetal and postnatal hearts, which are dominated by the thickness of the compact myocardium (Shi et al., 2023). What is entailed by alleged compaction varies between definitions (Wilsbacher and McNally, 2016). Most often, it is thought to involve the coalescence of trabeculations to produce the compact wall. The morphometric support for this process is weak (Faber et al., 2021a). The morphometric consequences of compaction, if they occurred, would be thickening of the compact layer by a reciprocal reduction of the trabecular layer, a reduction in the number of trabeculations, and a reduction in the number and size of the intertrabecular recesses (Faber et al., 2021a). Because the apical components of the human ventricles exhibit the highest trabecular-to-compact ratio, it is thought that compaction is least extensive in these regions, and most extensive at the ventricular base (Sedmera et al., 1999, Hussein et al., 2015, FinstererStollberger and Towbin, 2017).

Quantifications, such as those mentioned above, can in principle measure key aspects of compaction, as well as differences in the rate of growth of the trabecular and compact layers. For example, a decrease in the volume of trabecular myocardium could indicate compaction, whereas an increase in trabecular volumes would indicate growth. Compaction was initially quantified in chicken embryos. Specifically, the right ventricular free wall was assessed, which, in the adult bird, is very sparsely trabeculated, even when compared to the parietal wall of the neighboring left ventricle (Rychterova, 1971). Compared to many mammalian species, chicken is an extreme case simply because of these differences in trabeculation (Rowlatt, 1990). Indeed, the setting in human is the inverse, since the right ventricle is more trabeculated than is the left ventricle (Riekerk et al., 2022). Shrews, a group of tiny mammals, quite resemble chickens in the sparseness of trabeculation in the right ventricle, but the thickening of their compact wall during gestation occurs at a time when the trabecular layer itself is also increasing in width and volume (ChangSheftel and Jensen, 2022). Compaction, then, can play a role in cardiogenesis (Hanemaaijer et al., 2019). The magnitude of the process, however, is much more ambiguous when considering the extent to which it applies to various mammalian species, particularly human. Perhaps the most used measure of compaction is the trabecular-to-compact ratio. This measurement is itself a challenge. Should the value be interpreted as relating to the width of the trabecular layer as the numerator, the compact layer as the denominator, or both? This challenge is particularly acute concerning developmental studies, since the ratio changes dramatically during the continuous growth of the trabecular and compact layers, both in humans (BlausenJohannes and Hutchins, 1990, Faber et al., 2022b) and in mice (Ishiwata et al., 2003).

Here we used a comparatively large dataset of high resolution images of mouse hearts, supplemented with data from human hearts, to quantify the changes in the extent of the trabecular and compact layers that coincide with the proportional reduction of the trabecular layer that is interpreted as compaction. We show that the changes interpreted as compaction are found during a period of relatively slow growth of the trabecular layer.

## Materials and methods

### Specimens

We have studied over 450 datasets prepared by high-resolution episcopic microscopy showing the structure of the developing heart in wild-type mice of an outbred background (NIMR Parkes) from embryonic day (E) 9.5 to postnatal day 3. From these samples, we chose appropriate datasets for each of the days of development. The mice had been collected as part of an extensive study of normal and abnormal cardiac development undertaken initially at the National Institute of Medical Research, and continued subsequent to the transfer to the Crick Institute in London. Human embryos (Carnegie stages 11 – 20) and one 11 post conception week fetus were obtained from the MRC/Wellcome-Trust funded Human Developmental Biology Resource maintained at Newcastle University (www.hdbr.org). All the samples were imaged using high-resolution episcopic microscopy and micro-computed tomography techniques as previously described (GeyerMohun and Weninger, 2009, Degenhardt et al., 2010, Anderson and Bamforth, 2022). Briefly, stacks of intrinsically aligned serial images were subsampled and converted into volume data sets and analysed using Amira software (ThermoFisher Scientific). Three-dimensional images were created by manual segmentation using the label field function of Amira.

### Micro-computed tomography

The hearts of six adult mice (15 months) of either sex were perfused in Langendorff mode for the purposes of optical mapping of electrical impulse propagation (Olejnickova et al., 2021) followed by fixation for 24 h in 4% paraformaldehyde in PBS at 4 C. The hearts were then processed for micro-computed tomographic (CT) examination essentially as described recently (Gregorovicova et al., 2022), keeping the specimens in iodine solution for 1 month. The specimens were scanned in plastic tube immersed in phosphate-buffered saline with the following scanning parameters: pixel size = 7.5 μm, source voltage = 90 kV, source current = 111 μA, filter: Al 0.5mm + Cu 0.038mm, rotation step = 0.2°, frame averaging = 2, specimen rotation of 180°, camera binning 2x2, scanning time = approx. 3 hours per specimen. Flat-field correction was updated prior to each scanning. Scans were acquired using SkyScan 1272 (Bruker micro-CT, Belgium). Projection images were reconstructed with NRecon (Bruker micro-CT, Belgium) with the adequate setting of correction parameters (misalignment, smoothing, ring-artifact correction and beam hardening). 3D visualization was created by CT Vox (Bruker micro-CT, Belgium). CTAn (Bruker micro-CT, Belgium) was used to perform image processing.

### Quantifications

Image stacks were imported to Amira (v2020.2, Thermo-Fisher). On equidistant images, either in the transverse or frontal plane of the heart, we labeled the trabeculations as opposed to the compact layer of the ventricular walls for both left and right ventricles. Approximately 10 images were labeled for each heart (Figure 1). The hearts were generally clear of blood coagulates and the full depth of most intertrabecular recesses could be seen. In this way, the intertrabecular recesses mark the trabecular layer as distinct from the compact layer which is free of recesses (Figure 1). The labeled areas relate to the true volume, according to Cavalieri’s principle (Gundersen et al., 1988) or Simpson’s rule, with an error of approximately 10 %. Volume readouts of the labels were derived from the Materials Statistics module in Amira. They were then multiplied by the fixed distance between the labeled images. Especially the embryonic hearts were quite uniform in appearance, whereas the oldest hearts could have somewhat folded walls, but it has previously been shown that only gross mislabeling can substantially change the growth trajectories of fast growing myocardium (Faber et al., 2022b). The correlation between trabecular and compact volume was calculated with a linear regression analysis in Excel 2016 (Microsoft).

**Figure 1.**
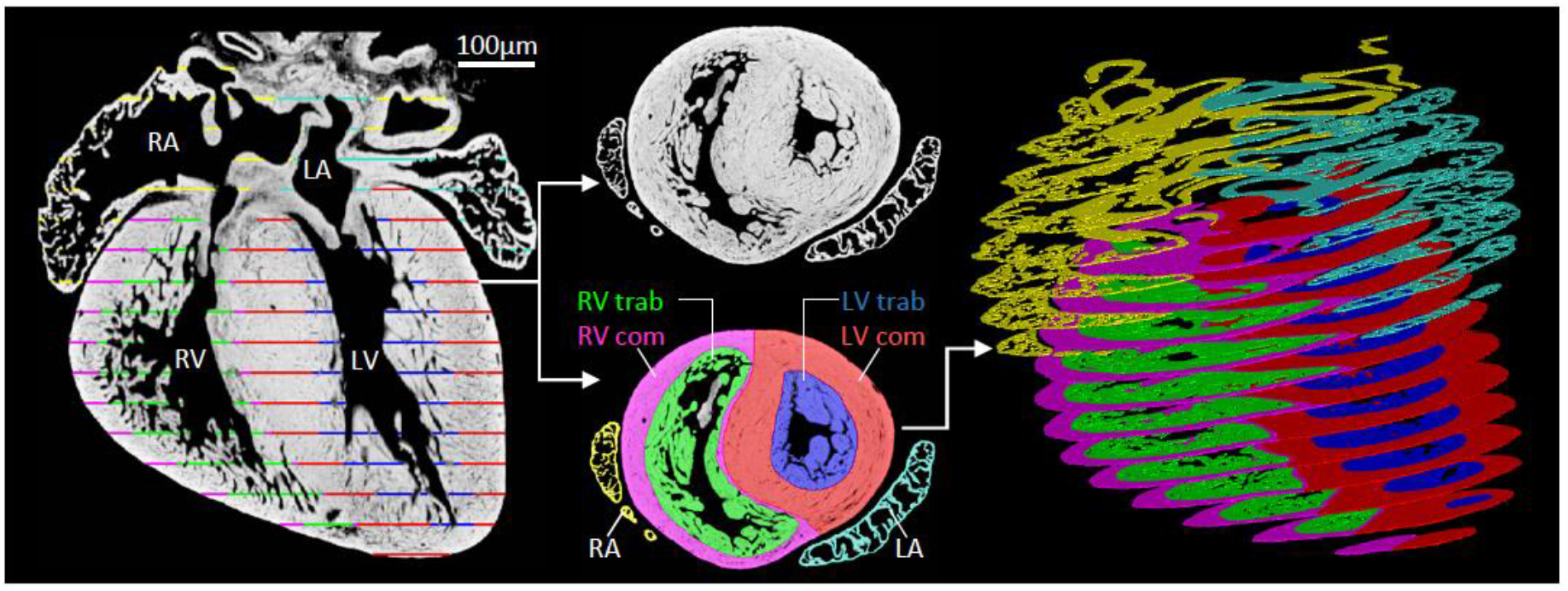
Work flow of myocardial volume quantifications. Approximately 10 equidistant images were chosen (left image) for labeling of trabecular (trab) and compact (com) myocardium of the left ventricle (LV) and right ventricle (RV). LA, left atrium; RA, right atrium.

So as to provide images, episcopic datasets were acquired for each day of development from embryonic (E) day 10.5 to gestation, which for the mouse is equivalent to E18.5. An additional dataset was examined from the third postnatal day. For each dataset, orthogonal sections were prepared in the frontal, sagittal, and short axis planes, as far as possible obtaining comparable sections for each dataset. The short axis sections originated from approximately mid-height in the apical region, where the apical region, in the fetal and postnatal hearts, can be defined as the part of the ventricle that is apical to the papillary muscles. Image stacks were imported using Osirix (Pixmeo SARL, Switzerland) software to create three-dimensional datasets, from which the images in orthogonal planes were selected.

## Results

### Quantifications of murine ventricular mural development

The volume of the trabecular and compact layers increases many fold from the smallest and youngest hearts, from 10.5 days since conception, to the ones obtained on neonatal day 3 (Figure 2). Compared to the left ventricle, the right ventricle attains a greater volume of trabeculations, and a smaller volume of compact muscle. Consequently, in the later stages, the right ventricle has a greater layer of trabecular myocardium compared to compact myocardium than does the left ventricle. In the early stages, the walls of both ventricles are composed primarily by trabecular myocardium, with three-quarters of the ventricular mass being occupied by trabeculations. By the third neonatal day, the trabeculations make up only about two-fifths of the volume of the right ventricular wall, but one-fifth of the volume of the left ventricular wall (Figure 2). This decrease in proportional trabeculation, however, is not brought about by a decrease in the volume of the trabeculations, as would be expected if the decrease was the outcome of coalescence of pre-existing trabeculations, or so-called “compaction”. Instead, the compact layer of the wall grows at a greater pace than does that of the trabeculations. Hence, it is the differential rate of growth that drives the reduction in proportional trabeculation (Figure 2).

**Figure 2.**
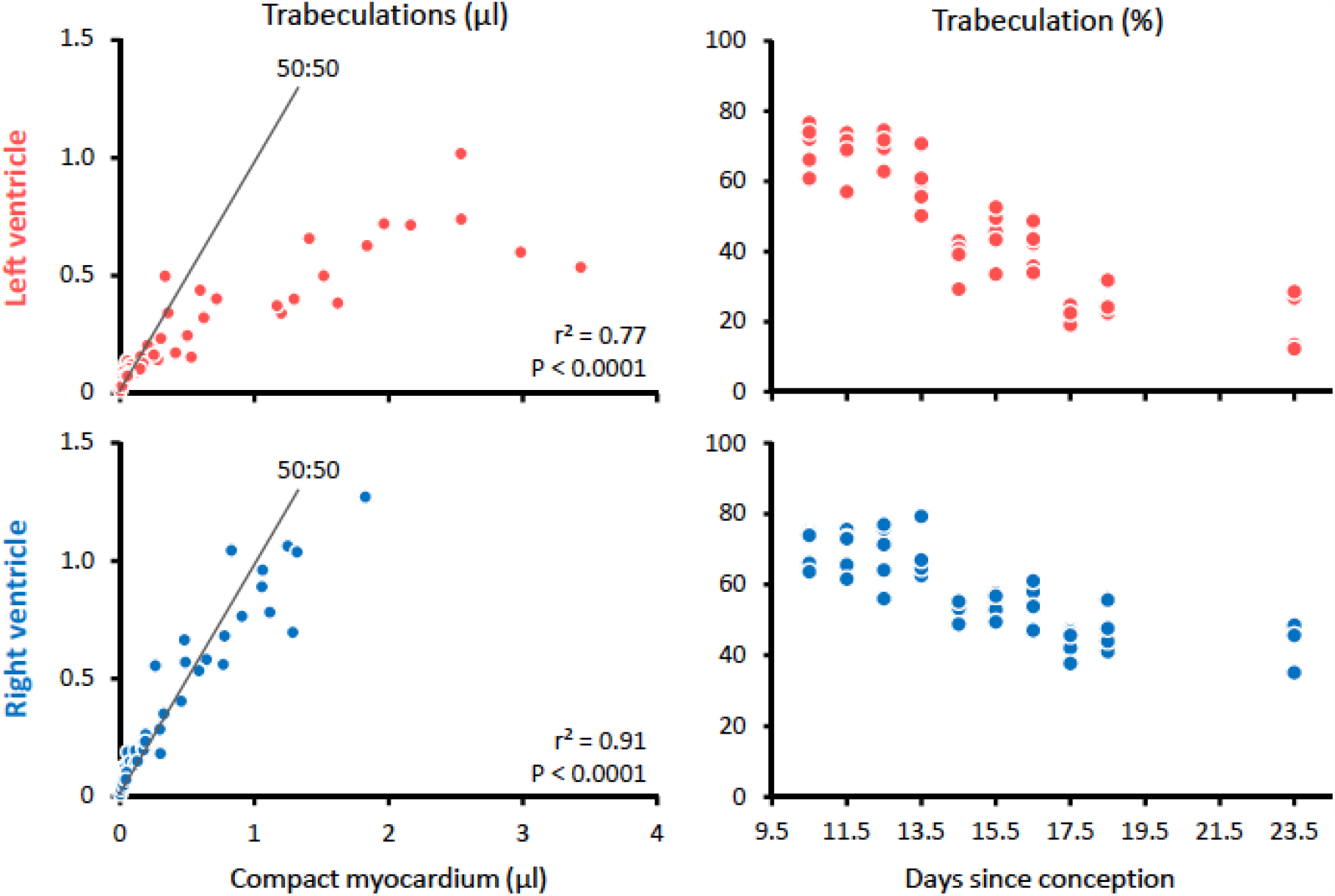
Change in trabecular volume and proportion in mouse heart development. Notice the strong positive and significant correlations between the volumes of compact and trabecular myocardium. This demonstrates that compaction, if measurable as a drop in trabecular volume and reciprocal increase in compact volume, does not drive growth of the compact wall. The line 50:50 indicates the relation at which trabecular and compact myocardium has the same proportion.

Having assessed the arrangement of the walls, we next quantified the volume of the intertrabecular recesses and central cavity of both ventricles of the one heart per period that had the median value of left ventricular trabeculations (Figure 3). When the volumes of the ventricular cavity were normalized to the volumes of ventricular myocardium, these ratios, for the early periods, were intermediate in value to those of end-diastolic and end-systolic left ventricles of adult humans (Figure 3B). In the fetal stages, the ratios decreased well below that of the end systolic value, suggesting that the fetal ventricles were much more contracted than the embryonic hearts (Figure 3B). Despite a likely greater state of contraction in the older and bigger hearts, we found a greater volume of the central cavity and intertrabecular recesses (Figure 3C). Proportionally, however, the intertrabecular recesses were smaller in the older hearts.

**Figure 3.**
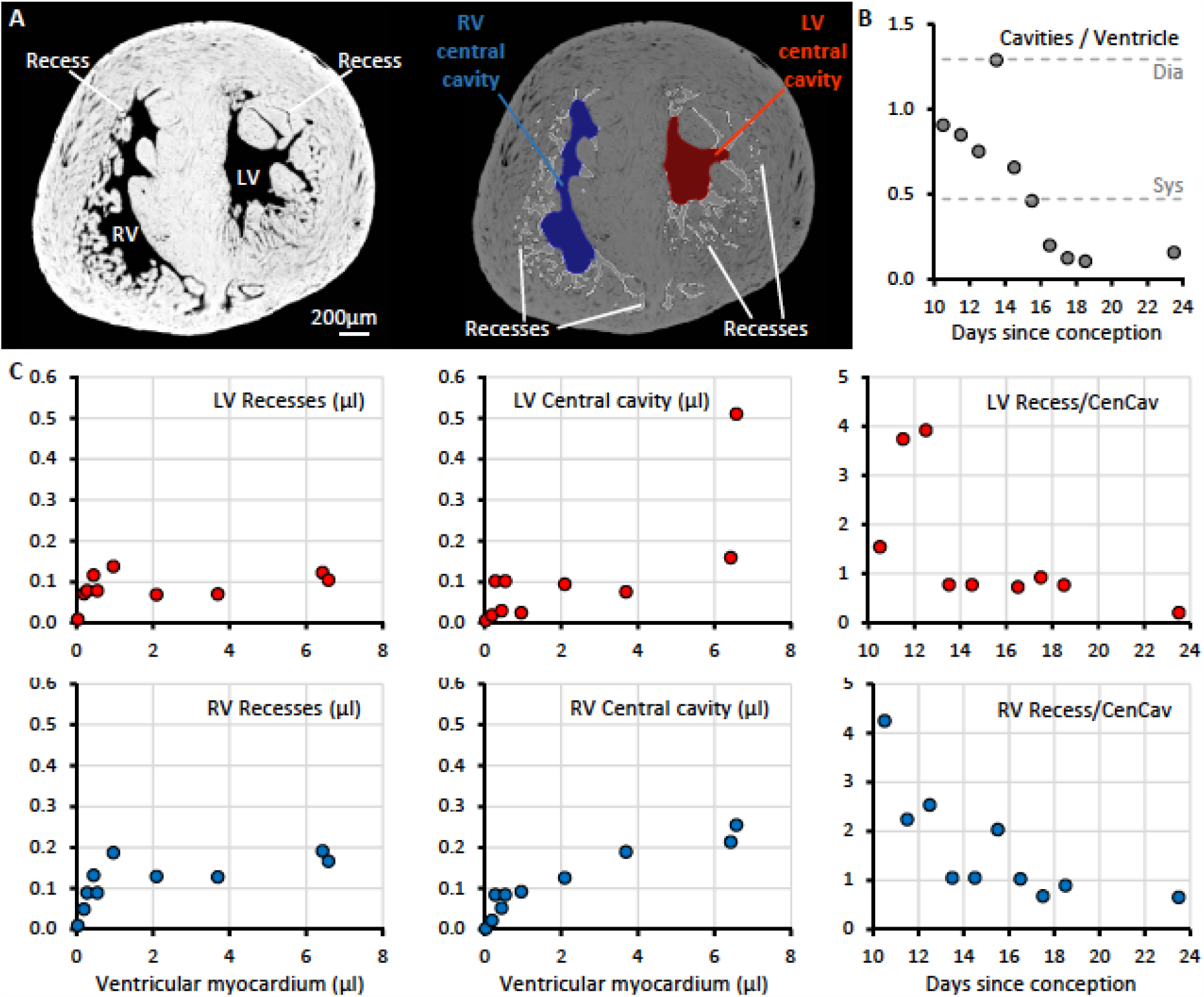
Gestational changes in the volumes of the ventricular cavities in mice. **A**. Short-axis mid-ventricular section of the E18.5 mouse ventricle, showing the labelling of central cavity and intertrabecular recesses. **B**. Lines Dia and Sys, refers to the ratio derived from dividing the adult human normal value of both sexes of left ventricular end diastolic volume (Dia) and end systolic volume (Sys) by the left ventricular mass (adapted from values from (Luu et al., 2022); The values are much the same in children (van der Ven et al., 2020)). In the mouse fetuses, the ratio of cavities to ventricular myocardial volume is well below the Sys line, suggesting these ventricles are much contracted. **C**. Gestational increase of the volume of the intertrabecular recesses and central cavity of both the left (red) and right ventricle (blue). There is a pronounced decrease in the proportion that the intertrabecular recesses comprise out of the total ventricular cavity.

Taken together, all myocardial and cavity volumes increase in development. Concomitantly, there is a decrease in the proportion that the trabeculations contribute to the ventricular walls. There is also a decrease in the proportion that the intertrabecular recesses contribute to the ventricular lumens. So as to provide a pictorial account of the changes, we chose, on the basis of the proportional changes, the median trabeculated specimen for each day of development so as to obtain the orthogonal planes through the left ventricle.

### Morphological description of murine ventricular wall development

Virtual sections were made for all five hearts for each of the periods from E10.5 through E18.5, and then comparable sections for the hearts obtained at the third postnatal day. Because of the pronounced changes in the anatomical arrangement of the ventricular walls during the periods from E13.5 to E15.5, it was difficult to provide comparable images for each specimen. Assessment of the datasets, however, showed that the specific plane of sectioning made little difference to the perceived thicknesses of the different layers. The hearts shown in Figures 4 through 6, therefore, are those with the median value of proportional left ventricular trabeculation for each day. As explained, for each dataset we prepared sections through the left ventricle in short axis, four chamber, and long axis planes.

**Figure 4.**
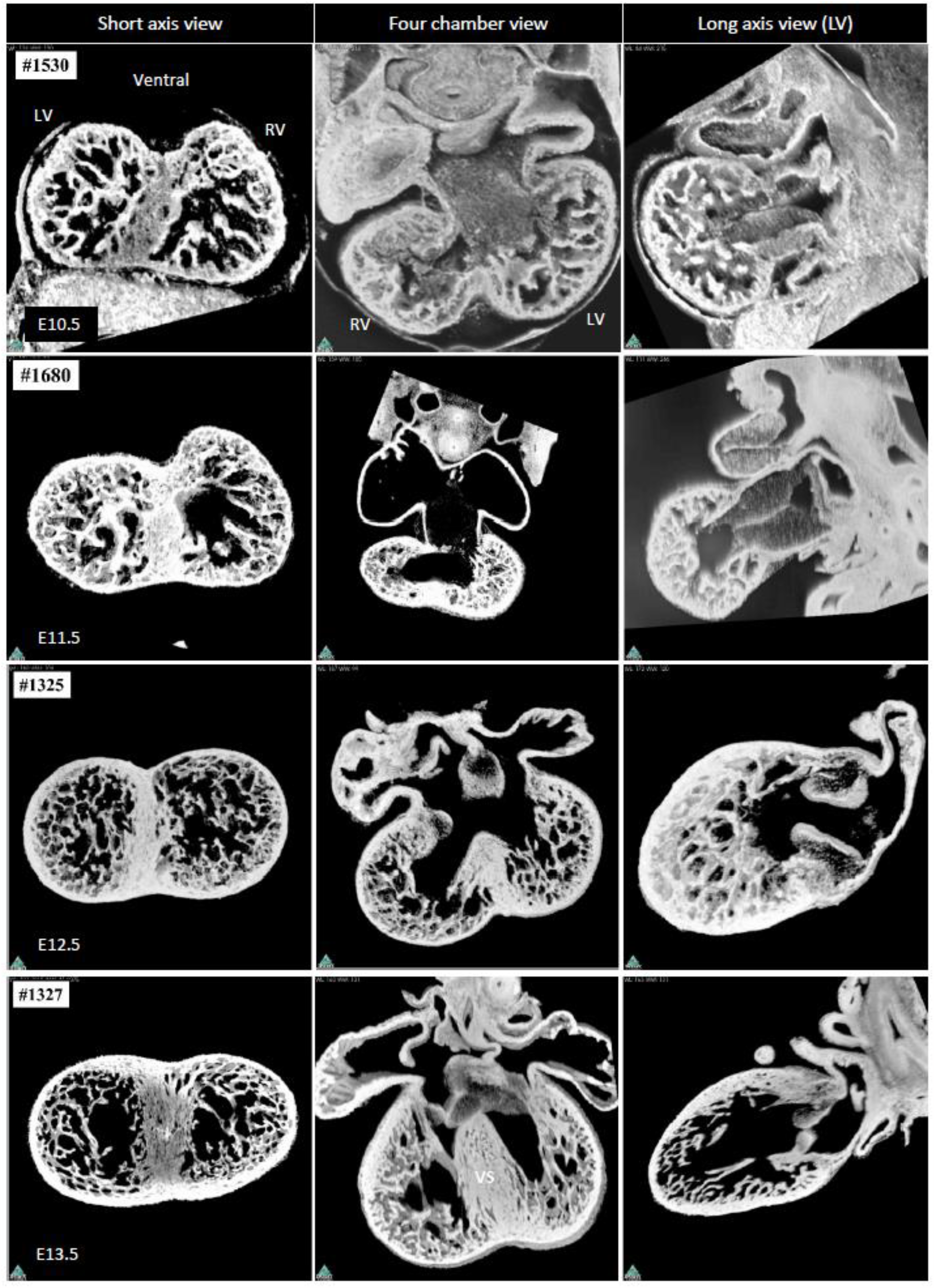
Embryonic development of the ventricular walls of mouse. LV, left ventricle; RV, right ventricle; VS, ventricular septum.

At the beginning of development, from E10.5 through E13.5, the ventricular walls are composed primarily of trabeculations (Figure 4). This coincides with the noted increase in trabecular mass (Figure 2). During the same period, nonetheless, we also observed a substantial increase in the volume of compact muscle (Figure 2). This change is most apparent when inspecting the ventricular septum (Figure 4, E13.5, label VS). It is during E13.5 that so-called “compaction” is alleged to contribute to the thickening of the compact wall (Faber et al., 2021a). Our data does not support that claim. A process of coalescence of trabeculations (Henderson and Anderson, 2009) is seen to occur so as to produce the papillary muscles of the atrioventricular valves, with a similar process producing the future septomarginal and septoparietal trabeculations of the right ventricle. None of this myocardium, however, is incorporated into the compact layer of the ventricular walls, but rather remains as trabeculated myocardium. The sides of the crest of the ventricular septum are also surmounted by trabeculations in the earliest stages. By E13.5, the septal crest is draped with only a thin layer of trabeculations. A significant portion of the initial trabeculations do not disappear, but instead become the branches of the atrioventricular conduction axis (van Weerd and Christoffels, 2016). Because of their low rate of growth, these surface trabeculations eventually become tiny when compared to the large bulk of the septum.

By E14.5, the central cavities of both ventricles are much expanded (Figures 4-5). This change has previously been measured as a decrease in the proportion that intertrabecular recesses compose of the total left ventricular cavity (Jensen et al., 2016). During this period, there is also a significant increase in the thickness of the compact layer itself. By E16.5 (Figure 5), the papillary muscles have become much thicker than the neighboring trabeculations. This finding shows that trabeculations themselves can grow well beyond their previous diminutive size without becoming incorporated into the compact layer of the wall. During the same stages, there is also a further “laying down” of the bundle branches on either side of the ventricular septum (Sankova et al., 2012). These changes are seen to continue during E17.5 (Figure 5). By this stage, when assessed in the short axis, the apical trabeculations are now no more than excrescences on the endothelial surface of the ventricular cavity, although when seen in the four chamber view, they provide a lace-like configuration in the right ventricle. By this stage, despite a high number of trabeculations, the compact layer has now increased markedly in thickness, and is also much greater in volume (Figure 2), with the proportion of trabecular muscle much lowered (Figure 2). During the period of transition from embryonic to fetal growth, therefore, major changes are seen towards adoption of the adult morphology, albeit without overt evidence of coalescence of trabeculations into the compact layer of the wall.

**Figure 5.**
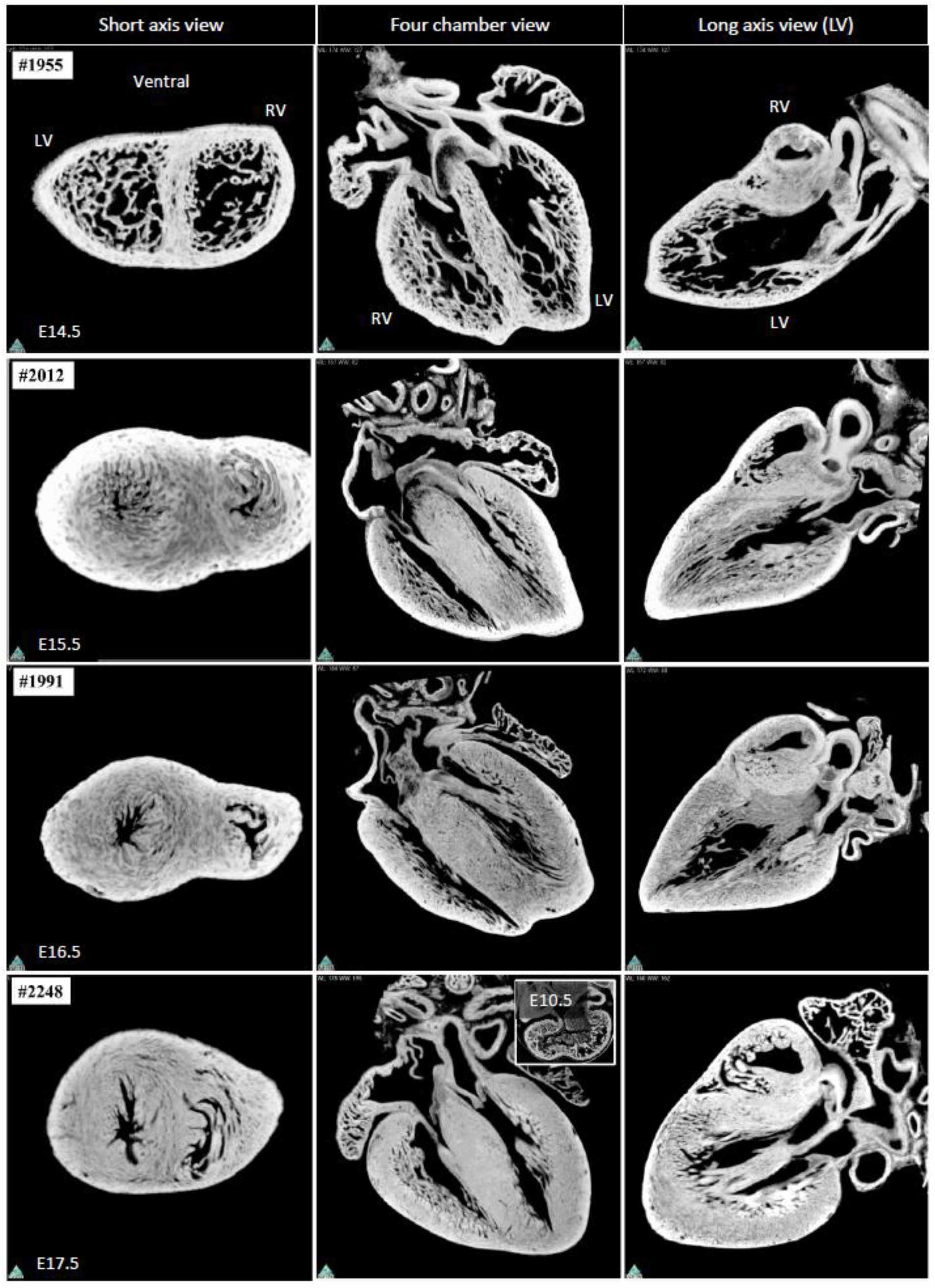
Fetal development of the ventricular walls of mouse. The panel with the 4-chamber view from the E17.5 period also contains the equivalent view of the heart of the E10.5 period, shown on the same absolute scale. LV, left ventricle; RV, right ventricle.

Towards the end of gestation, the compact layers of the ventricular walls begin to show aggregation of their contained cardiomyocytes into sheets (Figure 6). This remodeling is found exclusively within the compact wall, but does show some resemblance to a trabecular pattern. By the time of birth, the trabeculations are barely recognizable when compared to the thickness of the compact wall and the size of the ventricular cavity. The trabeculations, nonetheless, can still be recognized at the apex, with this feature seen in short axis and long axis cuts. They are particularly prominent beneath the bases of the papillary muscles, where they “cushion” the base of the muscles from the supporting compact layer of the walls. This perceived reduction in the trabecular layer is in part an effect of inspecting thin sections. Visualizations of the whole wall reveals a high number of trabeculations, even in the ventricular base which is considered to be the least trabeculated part of the ventricle (Figure 7).

**Figure 6.**
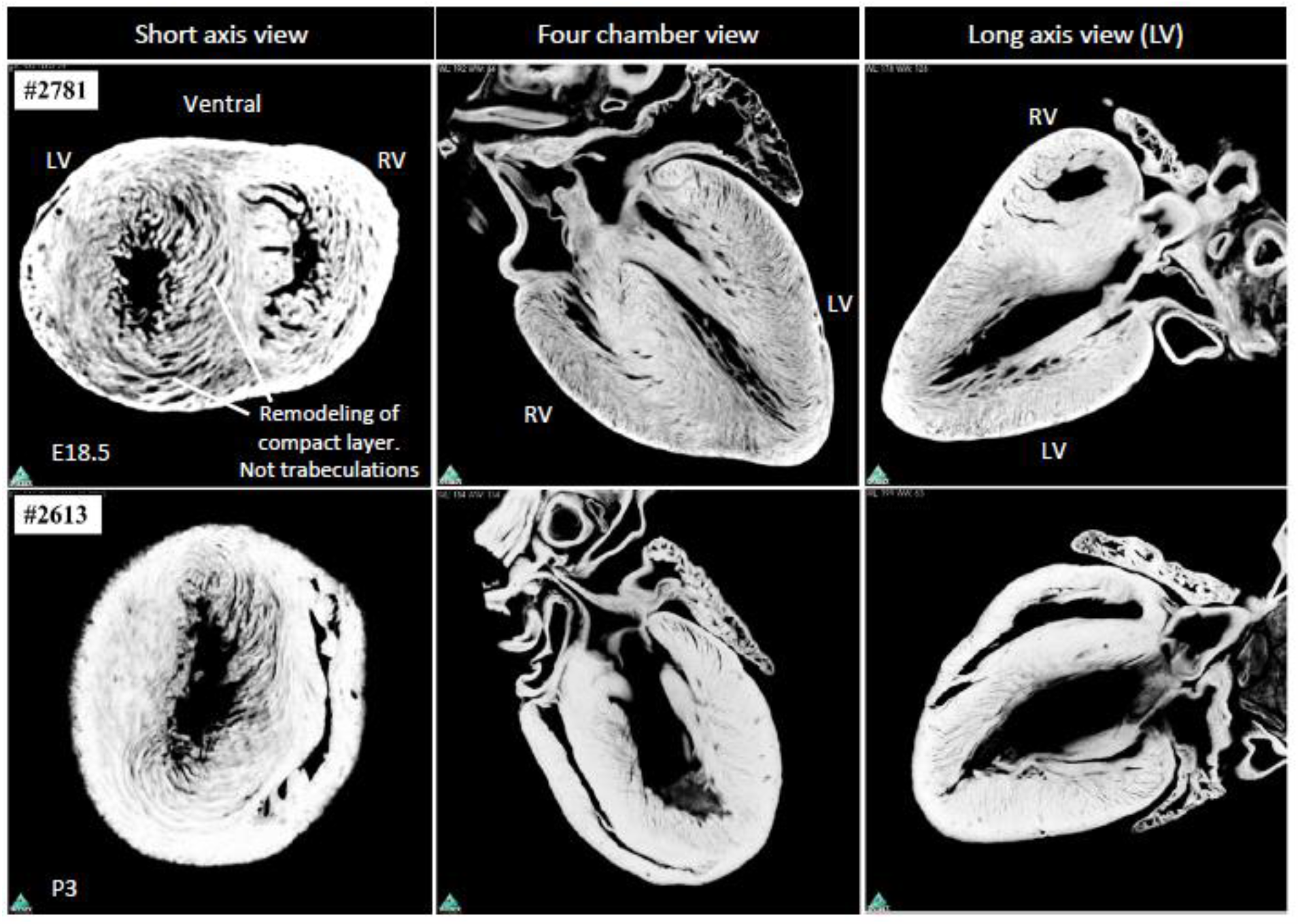
Perinatal development of the ventricular walls of mouse. LV, left ventricle; RV, right ventricle.

**Figure 7.**
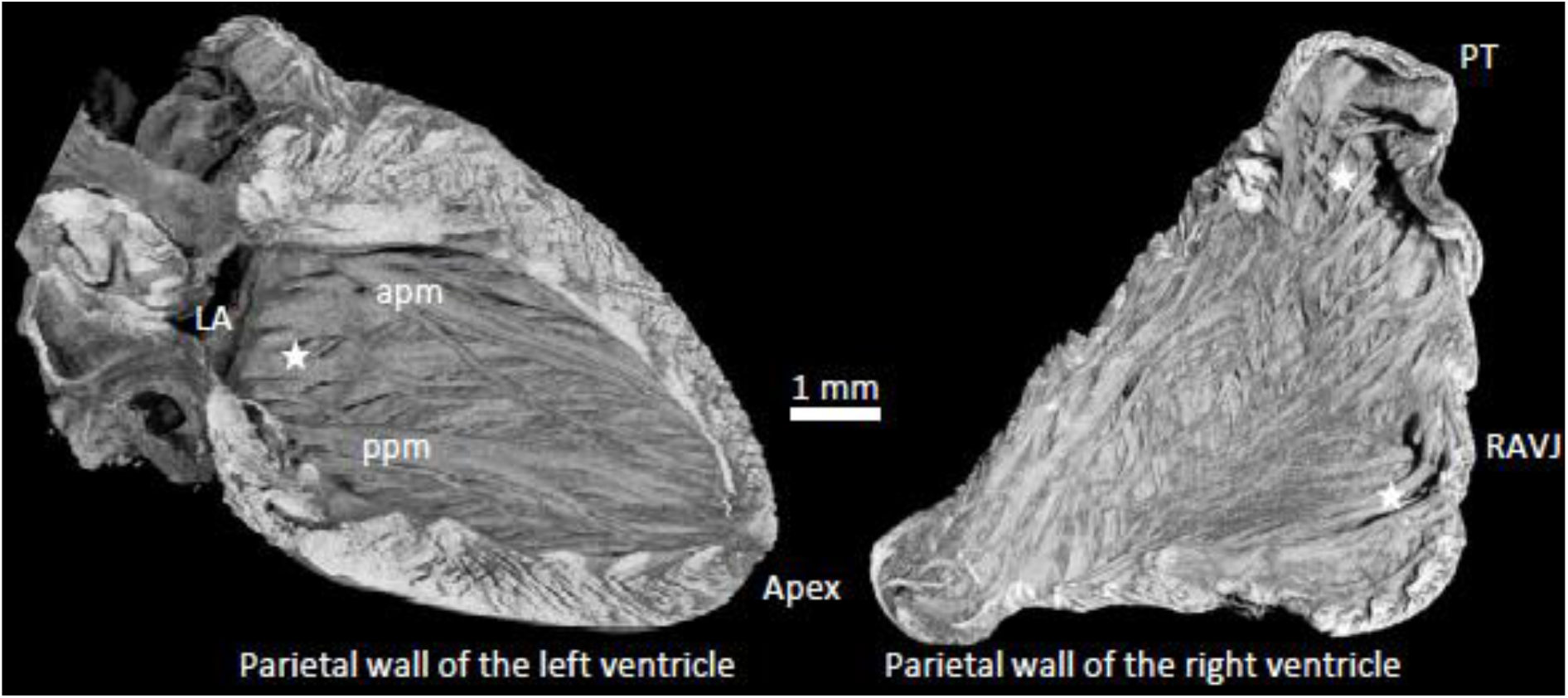
Extensive trabeculation in the adult mouse ventricles. The free wall of the adult mouse ventricles, reconstructed from micro-CT. While the ventricular base is considered the least trabeculated part of the ventricle, it nonetheless contains extensive trabeculation (see the area around the star). apm, anterior papillary muscle; LA, left atrium; ppm, posterior papillary muscle; PT, pulmonary trunk; RAVJ, right atrioventricular junction.

### Morphological description of human ventricular wall development

Figure 8 shows five periods of human gestation, from Carnegie stage 11 when the ventricular cavity is tiny, over a late embryonic stage at which time the ventricular septation is complete (Carnegie stage 20), to the early fetal period. In the early stages, from Carnegie stages 11 to 14, the ventricular walls are composed primarily by trabecular myocardium, which is visually apparent in Figure 8, and which has been quantified previously as a period of rapid growth (Faber et al., 2021a, Faber et al., 2021b). By Carnegie stage 20, the heart is much bigger, the compact wall is thicker, the central cavity has expanded, and the trabecular layer is proportionally reduced, whereas in absolute size the trabecular layer has grown substantially. For the 11^th^ week since conception, or the third week of fetal development, the heart shown in Figure 8 shows a pronounced, albeit not complete, morphological resemblance to the adult heart. Thus, the trabecular layer grows for every older stage, but not as much as the compact wall thickens and the central cavity expands.

**Figure 8.**
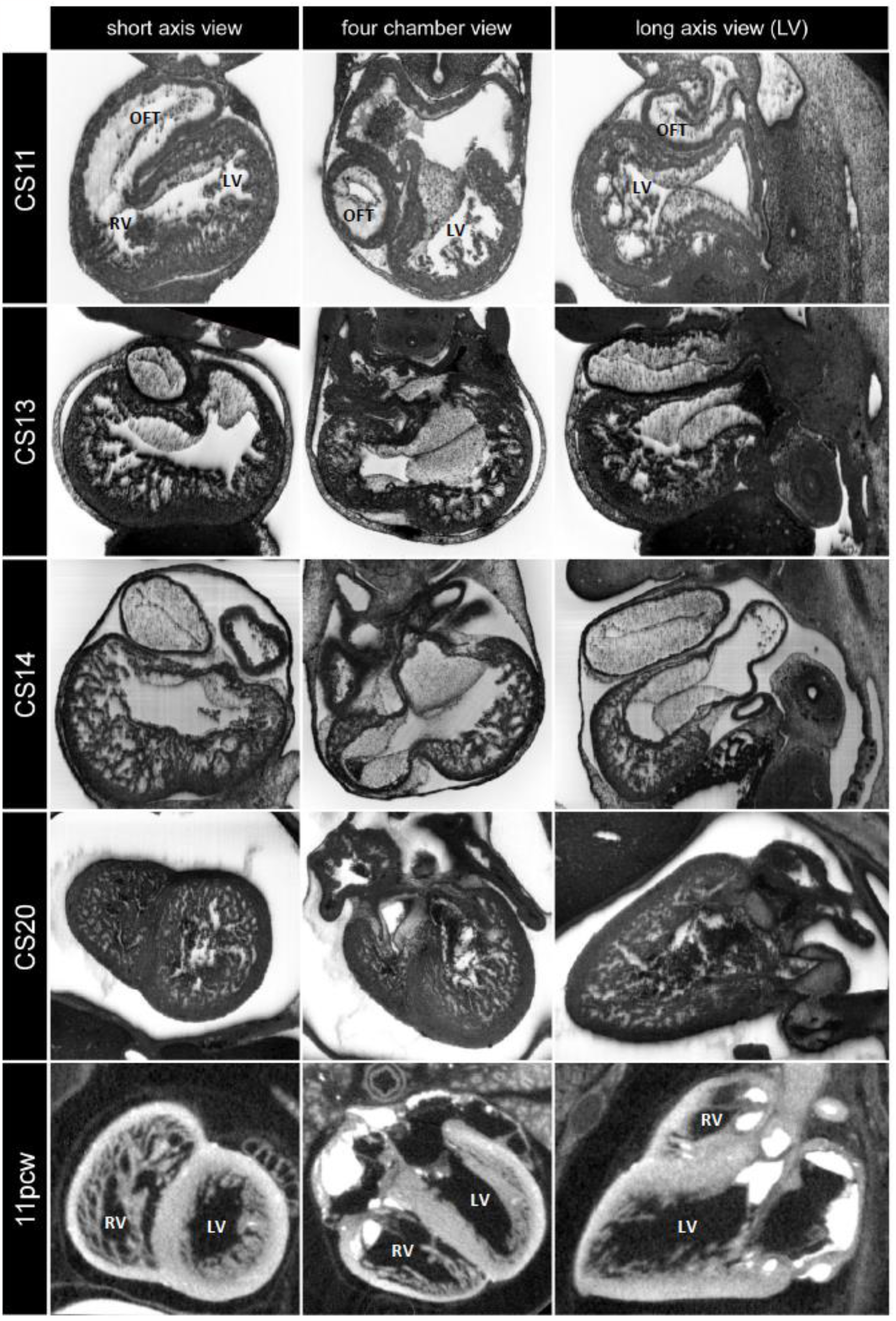
Embryonic and early fetal development of the ventricular walls of human. LV, left ventricle; OFT, outflow tract; RV, right ventricle.

## Discussion

Our data supports the notion that different rates of growth of the trabecular and compact layers is the major driver of the changes that take place in ventricular mural architecture. Compaction may seem to be an important factor if we look solely at the proportional reduction of the trabecular layer thickness. The interpretation of compaction as a major driver of morphological change, however, is confounded by several factors that occur simultaneously.

First among these is the absolute scale. The heart grows rapidly, and the number of cardiomyocytes increases from a few tens of thousands to approximately one million in the period compaction is said to occur (de Boer et al., 2012, Faber et al., 2021b). If images of fetal hearts are scaled down, and juxtaposed to microscopic embryonic hearts, e.g. (Oechslin and Jenni, 2011), this is in effect a reduction of the spatial resolution of the fetal hearts. So, while the fetal trabecular layer may seem to have flattened, in fact, the volume of trabeculations and intertrabecular recesses is greater in the fetal hearts. The impact of scale on spatial resolution, nonetheless, does not explain all the differences. Several factors are again in play. Some flattening of trabeculations likely occurs, most plausibly on the surface of the ventricular septum. This happens at a time when the ventricular cavities expand. The expansion may contribute to thin the trabecular layer while it is growing. This can be compared to making the base of a pizza from a lump of dough. With regard to the recesses, some may collapse and disappear, as suggested by lineage tracing of endocardium that gives rise to coronary endothelium (Tang et al., 2022). At the end point, however, the ventricle of the adult has such an extensive and intricate trabecular layer that it is exceedingly difficult to count the number of trabeculations or recesses, either in humans (Gerger et al., 2013, Riekerk et al., 2022) or in mice as we show in this report. While the extent of the trabecular layer is difficult to miss during *post mortem* macroscopic inspection, the degree of trabeculation is most frequently assessed during life by echocardiography and magnetic resonance imaging. These modalities, at present, do not have sufficient spatial resolution to show all trabeculations and recesses (Jensen and Petersen, 2022, Polacin et al., 2022, Riekerk et al., 2022).

The state of contraction is a second factor that affects the interpretation of the extent of the trabecular layer, also when embryos and fetuses are compared (Ishiwata et al., 2003). The embryonic ventricle will often appear to be in a somewhat diastolic state. Many fetal ventricles, however, will appear in a state of systole. Our own quantifications of the ratio of cavity-to-myocardial volume support this interpretation. If the fetal trabecular layer is relatively compressed, the intertrabecular recesses may appear to have diminished in number, if not disappeared. Previously, mouse hearts were perfusion-fixed under known preload pressure. It was found that both the trabecular and compact layers grew throughout development (Ishiwata et al., 2003). The importance of state of contraction is similarly well recognized when using clinical techniques, such as magnetic resonance imaging. The border between the trabecular and compact layers is relatively easy to identify in diastole, whereas in systole the intertrabecular recesses can become so compressed that part of the trabecular layer can be mistaken for compact wall (Grothoff et al., 2012). In our assessment, a greater state of contraction likely biases the transmural appearance towards one that suggests extensive compaction.

The third factor worthy of emphasis is the great changes that happen to the structures and spaces that surround the trabecular layer. It is during the period of greatest proportional decline of the trabecular layer (E13.5-17.5 in our mouse data sets), that the compact wall grows most rapidly, and the central cavity expands. From a point of view of cardiac output, rapid growth of the ventricular wall is unsurprising, since the rapidly growing embryo and fetus at that period requires a matching cardiac output. Soon after beating has commenced, heart rate is mostly constant, at around 150-200 beats per minute. Consequently, cardiac output in the growing embryo and fetus is sustained by an increase of the stroke volume. The ventricular cavities must expand accordingly (Faber et al., 2022a). The ventricular walls must then grow to maintain systolic pressure, in accordance with the law of Laplace (BuffintonFaas and Sedmera, 2013). In this regard, the systolic blood pressure is known to increase with development (Van Mierop and Bertuch, 1967, Ishiwata et al., 2003). In short, if the trabecular layer is assessed in proportion to the ventricular cavity and the compact wall, note must be taken of how much, and how fast, there is a shift in these baselines. It is plausible that the expansion of the central cavity goes a long way to explain how the trabeculations come to be laid down on the sides of the base of the ventricular septum.

A fourth factor for emphasis is that trabeculations can themselves coalesce to become fewer, yet of greater size, without impacting on the dimensions of the compact wall (Henderson and Anderson, 2009). Or, at the risk of creating an oxymoron, there can be compaction within the trabecular layer. Papillary muscles are the prime example of this process. From the beginning of development, trabeculations extend between the atrioventricular cushions, from which the leaflets will develop, and the compact wall (de Lange et al., 2004). In the adult hearts of humans and mice, the papillary muscles are between the valve leaflets and their trabecular base (Axel, 2004). While a reduction in the number of trabeculations and intertrabecular recesses is consistent with compaction, the development of the papillary muscle exemplifies that the same morphometric readout is equally consistent with coalescence of trabeculations within the trabecular layer. Extensive coalescence of trabeculations within the trabecular layer may occur in pig, since the proportional volumes of the trabecular layers are similar between pig and human, while the porcine walls have fewer trabeculations (Jensen et al., 2023).

Besides the confounding factors in assessing the extensiveness of the trabecular layer, in microscopy there is also a risk of mistaking compact wall for trabeculations. This relates to the fact that the cardiomyocytes of the compact wall become aggregated to form a three-dimensional meshwork during the later stages of fetal gestation (around E17.5-18.5 in our data) and such aggregates can resemble trabeculations on histology. Such myocardial organization likely allows for the aggregates to slide with respect to each other, thus allowing for a greater compression of the compact wall (Smerup et al., 2009). The risk of mistaking such components of the three-dimensional intramural mesh for trabeculations is much reduced with an endocardial specific stain, such as with antibody detection of endomucin (Gifford et al., 2019).

## Conclusion

Compaction is a developmental process that can substantially alter the morphology of some structures in some species, such as the right ventricular free wall in chicken (Rychterova, 1971). When assessing the morphological change to the trabecular layers of mice and humans, while a process of compaction may seem to provide an intuitive explanation for the thickening of the compact layer, other major processes are at play in parallel. The expansion of the central cavity is prominent among these, together with the intrinsic growth of the compact layer, the change of scale reflecting the tremendous and rapid overall growth of the heart, the coalescence of trabeculations within the trabecular layer itself, and the state of contraction of the walls. We posit that the true magnitude of compaction can only be revealed when account has been taken of these shifting baselines. For ventricular walls that contains a high number of trabeculations, such as the ones found in adult mouse and human, our findings indicate that compaction is unlikely to have played a major role in shaping their morphology.

## Acknowledgements

The authors have no conflicts of interest to declare.

## Notes

### Competing Interest Statement

The authors have declared no competing interest.

